# Oxycodone decreases anxiety-like behavior in the elevated plus-maze test in male and female rats

**DOI:** 10.1101/2021.12.02.470973

**Authors:** Adriaan W. Bruijnzeel, Azin Behnood-Rod, Wendi Malphurs, Ranjithkumar Chellian, Robert M. Caudle, Marcelo Febo, Barry Setlow, John K. Neubert

## Abstract

The prescription opioid oxycodone is widely used for the treatment of pain in humans. Oxycodone misuse is more common among people with an anxiety disorder than those without one. Therefore, oxycodone might be misused for its anxiolytic properties. We investigated if oxycodone affects anxiety-like behavior in adult male and female rats. The rats were treated with oxycodone (0.178, 0.32, 0.56, or 1 mg/kg), and anxiety-like behavior was investigated in the elevated plus-maze test. Immediately after the elevated plus-maze test, a small open field test was conducted to determine the effects of oxycodone on locomotor activity. In the elevated plus-maze test, oxycodone increased the percentage of time spent on the open arms, the percentage of open arm entries, time on the open arms, open arm entries, and the distance traveled. The males treated with vehicle had a lower percentage of open arm entries than the females treated with vehicle, and oxycodone treatment led to a greater increase in the percentage of open arm entries in the males than females. Furthermore, the females spent more time on the open arms, made more open arm entries, spent less time in the closed arms, and traveled a greater distance than the males. In the small open field test, treatment with oxycodone did not affect locomotor activity or rearing. Sex differences were observed; the females traveled a greater distance and displayed more rearing than the males. In conclusion, oxycodone decreases anxiety-like behavior in rats, and oxycodone has a greater anxiolytic-like effect in males than females.

## 1. INTRODUCTION

Prescription opioids are widely used for the treatment of chronic pain, and in 2015 more than one-third of adults in the United States used prescription opioids (Han et al., 2017). Thirteen percent of the adults with an opioid prescription misuse opioids (Han et al., 2017). During the last two decades, there has been a strong increase in drug overdose deaths. The number of overdose death increased from fewer than 20,000 in 1999 to more than 70,000 in 2019 (Centers for Disease Control and Prevention, 2021). The majority of overdose deaths are due to opioids (Seth et al., 2018), and it has been projected that opioid overdose deaths could reach more than 80 thousand by 2025 (Chen et al., 2019). Therefore, it is critical to understand what drives the misuse of prescription opioids.

Although opioids are mainly prescribed for pain, there is a strong association between prescription opioid use and psychiatric disorders. In particular, the use of prescription opioids is common among people with mood and anxiety disorders (Martins et al., 2012; Sullivan et al., 2006). Almost 50 percent of individuals who are dependent on prescription opioids suffer from a mood or anxiety disorder (Gros et al., 2013). People with anxiety disorders are more likely to start using prescription opioids, and the use of prescription opioids is more likely to lead to the development of dependence in people with anxiety disorders (Martins et al., 2009a). A study with chronic pain patients showed that those with an anxiety disorder were five times more likely to misuse their opioid medication than those without an anxiety disorder (Feingold et al., 2017). Interestingly, oxycodone decreases anxiety in patients with cancer pain (Sui-Hui et al., 2013). Therefore, it might be possible that prescription opioids are misused because they decrease anxiety. Indeed, adolescents report using opioids to self-medicate anxiety symptoms (Boyd et al., 2006). The misuse of prescription opioids is of concern because of the addictive properties of opioids and the highly aversive withdrawal signs that drive continued opioid use (Bruijnzeel et al., 2006; Srivastava et al., 2020).

Oxycodone is one of the most widely prescribed opioids for pain (Kane, 2021). Oxycodone is a semisynthetic opioid derived from the opium alkaloid thebaine. Oxycodone is considered a strong analgesic and has a high oral bioavailability compared to morphine (Riley et al., 2008). Oxycodone activates mu, delta, and kappa 2b-opioid receptors (Nielsen et al., 2007; Yang et al., 2016) and has a higher affinity for mu-opioid receptors (Ki value, 18 nM) than for delta (Ki value, 958 nm) and kappa-opioid receptors (Ki value, 677 nm)(Monory et al., 1999). Animal studies show that oxycodone has analgesic and rewarding properties (Collins et al., 2016; Liu et al., 2009; Nielsen et al., 2007; Pradhan et al., 2010).

Clinical studies indicate that prescription opioid abuse is common in people with anxiety disorders (Boyd et al., 2006; Martins et al., 2009a; Zacny et al., 2011). Even though oxycodone is widely prescribed in the United States and is more likely to be misused by people with an anxiety disorder than those without (Kane, 2021; Martins et al., 2009b), little is known about its effects on anxiety. Furthermore, despite extensive evidence for sex differences in the behavioral effects of opioids, little is known about sex differences in the effects of oxycodone in anxiety tests (Craft, 2008). Some studies have investigated the effects of selective mu-opioid receptor agonists on anxiety-like behavior. Treatment with the mu-opioid receptor agonist DAMGO ([D-Ala^2^,N-Me-Phe^4^-Gly^5^-ol]-enkephalin) does not affect anxiety-like behavior in the elevated plus-maze test in rats (Alexeeva et al., 2012); however, there is evidence that treatment with the opioid drugs morphine and fentanyl decreases anxiety-like behavior (Fujii et al., 2019; Koks et al., 1999; Rezayof et al., 2009). The goal of the present study was to investigate the effects of acute oxycodone administration on anxiety-like behavior and locomotor activity in adult male and female rats. Anxiety-like behavior was determined in the elevated plus-maze test and locomotor activity in the small open field test.

## 2. MATERIALS AND METHODS

### 2.1. Animals

Male and female Sprague Dawley rats (50 males and 50 females, 200-225 g) were purchased from Charles River (Raleigh, NC). The rats were kept in standard housing conditions (2 rats of the same sex per cage) in a climate-controlled vivarium on a reversed 12 h light-dark cycle (light off at 7 AM). Food and water were available ad libitum in the home cage. The experimental protocols were approved by the University of Florida Institutional Animal Care and Use Committee.

### 2.2. Drugs

Oxycodone hydrochloride was obtained from the NIDA Drug Supply Program and dissolved in sterile phosphate-buffered saline. The oxycodone solution was administered intraperitoneally (IP) in a volume of 1 ml/kg body weight.

### 2.3. Experimental design

The rats received injections with oxycodone (0, 0.178, 0.32, 0.56, and 1 mg/kg, IP) and were placed in the elevated plus-maze 30 min later. Immediately after the rats were tested in the elevated plus-maze test (40 min post-injection), the rats were placed in the small open field. Both the elevated plus-maze test and the small open field test lasted 5 min.

### 2.4. Behavioral tests

#### 2.4.1. Elevated plus maze test

The elevated plus-maze test is used to measure anxiety-like behavior in rodents (Walf and Frye, 2007) and was performed as described in our previous work (Chellian et al., 2020; Qi et al., 2016; Tan et al., 2019). The elevated plus maze (Coulbourn Instruments, Holliston, MA) consists of two open arms (i.e., without walls; 50 cm × 10 cm; L × W) and two closed arms (i.e., with black walls, 50 cm × 10 cm × 30 cm; L × W × H). The open and closed arms were connected by a central platform, and the open arms had 0.5 cm tall ledges to prevent the rats from falling off. The open arms were placed opposite of each other, and the maze was elevated 55 cm above the floor on acrylic legs. At the beginning of each test, the rats were placed in the central area facing an open arm and were allowed to explore the apparatus for 5 min. The rats were recorded with a camera mounted above the maze. The test was conducted in a quiet, dimly lit room (100 lux). The open arm and closed arm duration, number of open and closed arm entries, and total distance traveled were determined automatically (center-point detection) using EthoVision XT 11.5 software (Noldus Information Technology, Leesburg, VA). The percentage of open arm entries (open arm entries/total arm entries) and percentage time on the open arms (open arm time/total time on the arms) were calculated. Heatmaps were produced with the EthoVision heatmap generator. The apparatus was cleaned with a Nolvasan solution between tests.

#### 2.4.2. Small open field test

The small open field test was done to assess locomotor activity, rearing, and stereotypies (Bruijnzeel et al., 2016; Qi et al., 2016). Horizontal and vertical beam breaks were measured using an automated animal activity cage system (VersaMax Animal Activity Monitoring System, AccuScan Instruments, Columbus, OH, USA). Horizontal beam breaks and total distance traveled reflect locomotor activity, and vertical beam breaks reflect rearing. Repeated interruptions of the same beam is an estimate of stereotypies (Chellian et al., 2020; Febo et al., 2003; Hayashi et al., 2007). The setup consisted of four animal activity cages made of clear acrylic (40 cm × 40 cm × 30 cm; L x W x H), with 16 equally spaced (2.5 cm) infrared beams across the length and width of the cage. The beams were located 2 cm above the cage floor (horizontal activity beams). An additional set of 16 infrared beams was located 14 cm above the cage floor (vertical activity beams). All beams were connected to a VersaMax analyzer, which sent information to a computer that displayed beam data through Windows-based software (VersaDat software). The small open field test was conducted in a dark room, and the cages were cleaned with a Nolvasan solution (chlorhexidine diacetate) between animals. Each rat was placed initially in the center of the small open field, and activity was measured for 5 min.

### 2.5. Statistics

Elevated plus-maze and small open field data were analyzed using a two-way ANOVA, with treatment condition (oxycodone dose) and sex as between-subject factors. For all statistical analyses, significant interactions in the ANOVA were followed by Bonferroni’s posthoc tests to determine which groups differed from each other. P-values that were less or equal to 0.05 were considered significant. Significant main effects, interaction effects, and posthoc comparisons are reported in the Results section. Data were analyzed with SPSS Statistics version 27 and GraphPad Prism version 9.2.0.

## 3. RESULTS

### 3.1. Oxycodone and the elevated plus-maze test

Oxycodone increased the percentage of time that the rats spent on the open arms (Treatment F4,90=4.133, P<0.01; Sex x Treatment F4,90=0.307, NS, Fig. 1A), and increased the percentage of open arm entries (Treatment F4,90=3.998, P<0.01, Fig. 1B). Treatment with oxycodone differently affected the percentage of open arm entries in the males and the females (Sex x Treatment F4,90=2.594, P<0.05). The posthoc showed that females treated with vehicle had a higher percentage of open arm entries than males treated with vehicle, and oxycodone induced a greater increase in the percentage of open arm entries in the males than the females. Treatment with oxycodone increased the time on the open arms (Treatment F4,90=5.029, P<0.01; Sex x Treatment F4,90=0.366, NS, Fig. 2A), but did not affect open arm entries (Treatment F4,90=2.127, NS; Sex x Treatment F4,90=1.117, NS, Fig. 2B). Oxycodone decreased the amount of time spent in the closed arms (Treatment F4,90=3.587, P<0.01; Sex x Treatment F4,90=0.361, NS, Fig. 2C), and did not affect closed arm entries (Treatment F4,90=0.329, NS; Sex x Treatment F4,90=0.291, NS, Fig. 2D). Furthermore, oxycodone increased the total distance traveled (Treatment F4,90=4.833, P<0.01; Sex x Treatment F4,90=1.421, NS, Fig. 2E).

**Figure 1.**
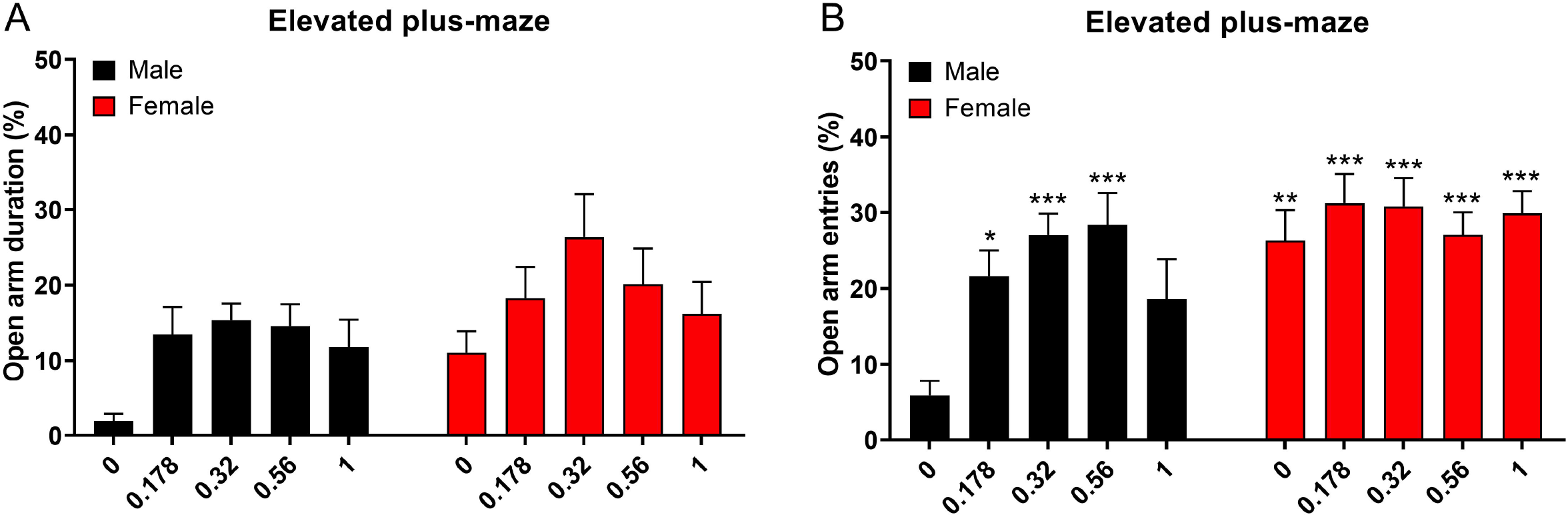
Oxycodone increases the percentage of open arm time and the percentage of open arm entries in the elevated plus-maze test. Oxycodone increased the percentage of time on the open arms (A) and the percentage of open arm entries (B) in the male and female rats. Compared to the males, the females spent a higher percentage of time on the open arms and had higher percentage open arm entries. Asterisks indicate a higher percentage of time on the open arms compared to the male rats treated with vehicle. *, p<0.05; **, p<0.01; ***, p<0.001. N=10/group. Data are expressed as means ± SEM.

**Figure 2.**
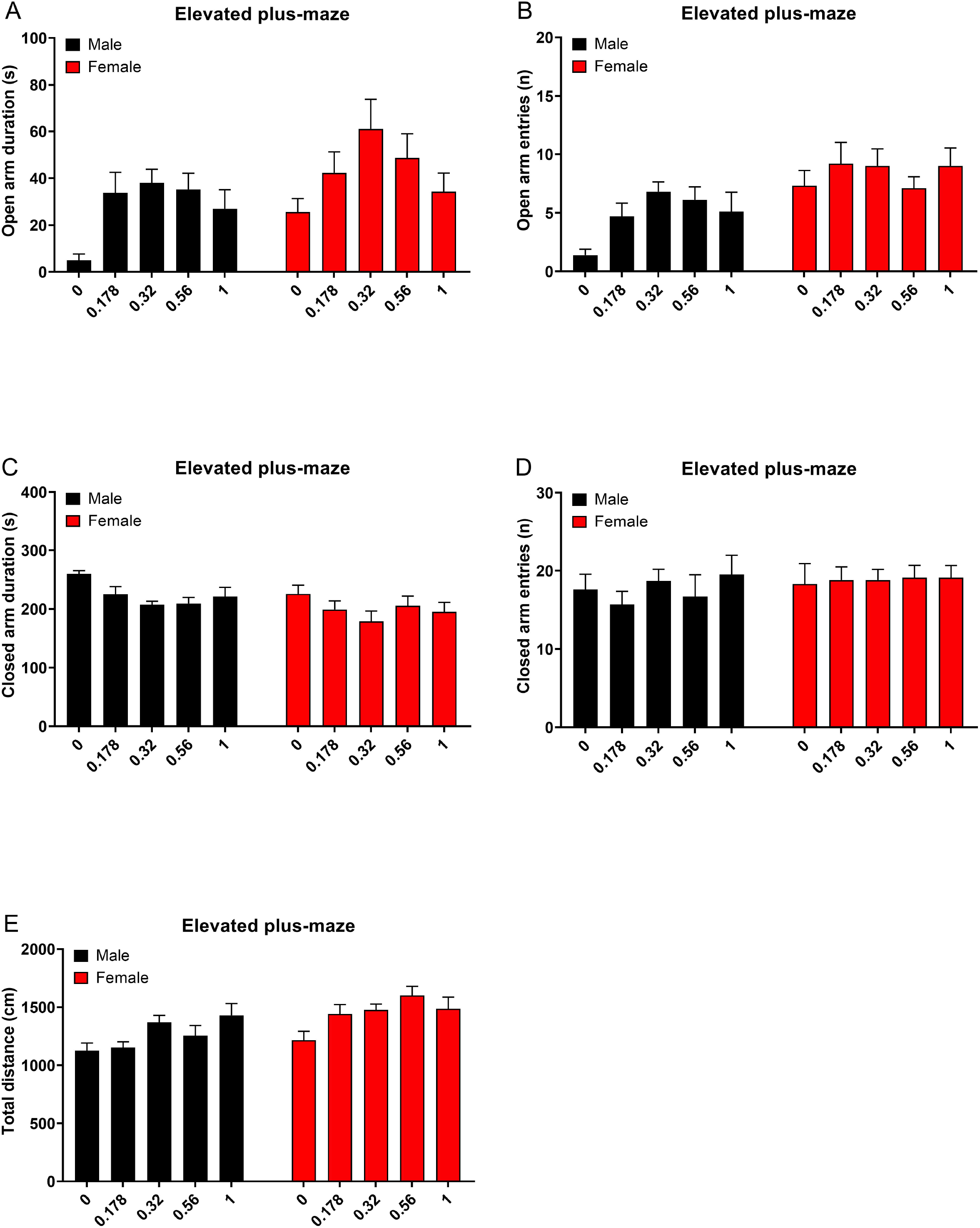
Oxycodone increases open arm time and decreases close arm time in the elevated plus-maze test. Oxycodone increased the time on the open arms (A) and did not affect open arm entries (B). Treatment with oxycodone decreased the amount of time spent in the closed arms (C) and did not affect closed arm entries (D). Oxycodone increased the total distance traveled (E). N=10/group. Data are expressed as means ± SEM.

There was a main effect of sex for almost all behavioral parameters. Compared to the males, the females spent a higher percentage of time on the open arms (Sex F1,90=8.853, P<0.01, Fig. 1A) and had a higher percentage of open arm entries (Sex F1,90=14.727, P<0.001, Fig. 1B). The females spent more time on the open arms (Sex F1,90=7.893, P<0.01, Fig. 2A) and made more open arm entries (Sex F1,90=19.418, P<0.001, Fig. 2B). The females also spent less time in the closed arms (Sex F1,90=7.521, P<0.01, Fig 2C) and traveled a greater distance compared to the males (Sex F1,90= 13.242, P<0.001 Fig 2E). The number of closed arm entries was not affected by the sex of the rats (Sex F1,90=0.894, NS; Fig 2D). Elevated plus-maze heat maps are presented in Figure 3.

**Figure 3.**
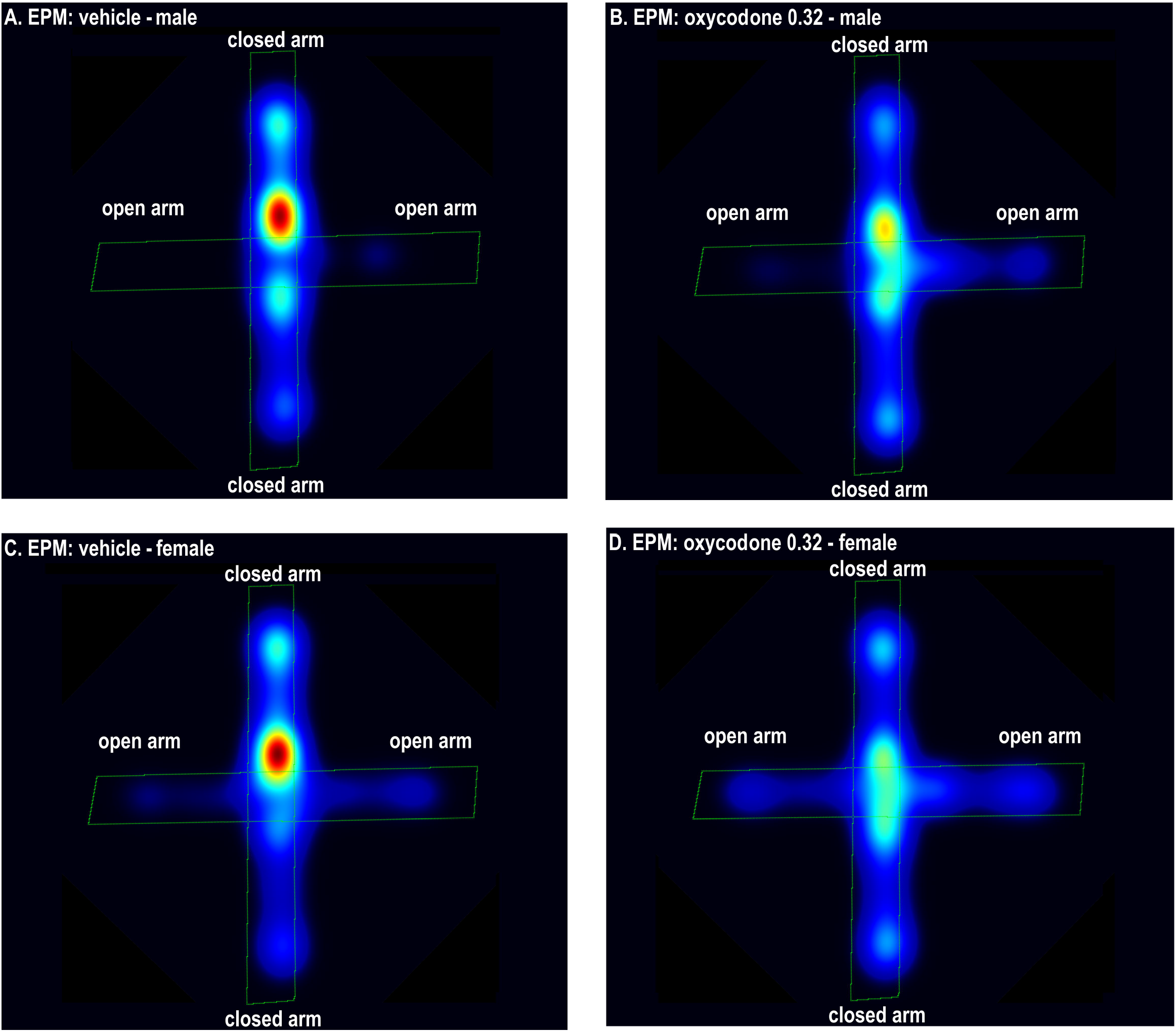
Heatmaps of the effects of oxycodone and sex in the elevated plus-maze test. The figures show that oxycodone (0.32 mg/kg) increased the amount of time on the open arms in the males (A, B) and the females (C, D). The females spent more time on the open arms than the males. The figures were generated with EthoVision software, and warmer colors indicate more time spent in a specific area. N = 10/group. Abbreviations: EPM, elevated plus-maze.

### 3.2. Oxycodone and the small open field test

Treatment with oxycodone did not affect the total distance traveled (Treatment F4,90=0.971, NS; Sex x Treatment F4,90=0.982, NS, Fig. 4A), horizontal beam breaks (Treatment F4,90=1.132, NS; Sex x Treatment F4,90=0.959, NS, Fig. 4B) or stereotypies (Treatment F4,90=0.783, NS; Sex x Treatment F4,90=0.632, NS, Fig. 4D). Treatment with oxycodone differently affected rearing in the males and the females (Treatment F4,90=0.97, NS; Sex x Treatment F4,90=2.738, P<0.05, Fig. 4C), but the posthoc comparison did not reveal any significant differences between the groups.

**Figure 4.**
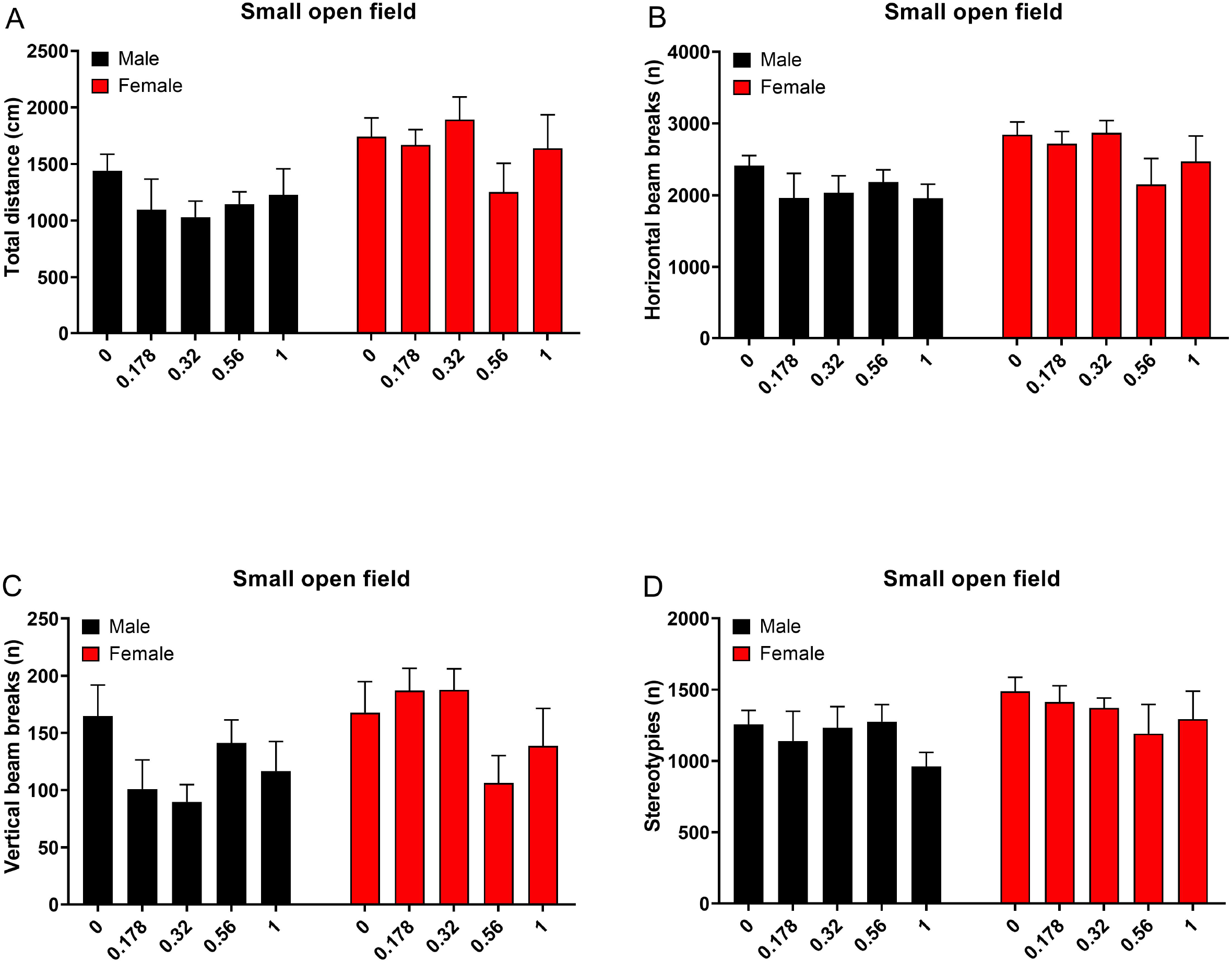
Oxycodone does not affect activity in the small open field test. Rats were treated with oxycodone and tested in the small open field 40 min later. Treatment with oxycodone did not affect the total distance traveled (A), horizontal beam breaks (B), vertical beam breaks (C), and stereotypies (D). The females traveled a greater distance and had more horizontal and vertical beam breaks than the males. N=10/group. Data are expressed as means ± SEM.

As in the elevated plus-maze, there was a main effect of sex for most behavioral parameters in the small open field test. Compared to the males, the females traveled a greater distance (Sex F1,90=12.144, P<0.01, Fig. 4A), and had more horizontal beam breaks (Sex F1,90=10.199, P<0.01, Fig. 4B) and vertical beam breaks (Sex F1,90=5.254, P<0.05 Fig. 4C). There was no sex difference in the number of stereotypies (Sex F1,90=3.798, NS Fig. 4D).

## 4. DISCUSSION

The goal of the present study was to investigate the effects of oxycodone on anxiety-like behavior in male and female rats. The results show that oxycodone increases the percentage of time on the open arms and the percentage of open arms entries in the elevated plus-maze test. Oxycodone did not affect closed arm entries in the elevated plus-maze test, suggesting that doses that decreased anxiety-like behavior did not have sedative effects. Furthermore, in the small open field test, oxycodone did not affect locomotor activity, rearing, or stereotypies. In the elevated plus-maze test, females had a higher percentage of open arm entries and spent a greater percentage of time on the open arms than the males. The females were also more active than the males in the small open field test. Considered together, these findings indicate that oxycodone reduces anxiety-like behavior in both sexes at doses that do not have sedative effects.

In the present study, we investigated the effects of oxycodone on anxiety-like behavior in the elevated plus-maze test. The elevated plus-maze has been widely used to investigate the effects of stressors and drugs on anxiety-like behavior. Stressors and anxiogenic drugs increase anxiety-like behavior in the elevated plus-maze test (Adamec et al., 1991; Korte et al., 1999). In contrast, anxiolytic drugs decrease anxiety-like behavior in the elevated plus-maze test. For example, the long-acting benzodiazepines diazepam and chlordiazepoxide increased the percentage of time on the open arms and the percentage of open arms entries (File, 1990; Guimarães et al., 1990; Stock et al., 2000). The present study shows that acute treatment with oxycodone has potent anxiolytic-like effects in the elevated plus-maze test. Oxycodone increased the percentage of time on the open arms and the percentage of open arm entries, which are some of the most widely used parameters to evaluate the effects of anxiolytic drug treatments (Hogg, 1996; Pellow and File, 1986).

In the present study, the rats that were treated with oxycodone were also evaluated in the small open field test. Treatment with oxycodone did not affect the total distance traveled, horizontal beam breaks, rearing, or stereotypies. This observation is in line with a prior study showing that low doses of oxycodone (0.25-1 mg/kg) do not affect locomotor activity, whereas high doses (2 mg/kg or higher) decrease locomotor activity in male and female Sprague Dawley rats (Holtman Jr and Wala, 2006). In our study, the rats were tested in the small open field 40 min after the administration of oxycodone. As the half-life of oxycodone in rats is 40-60 min (Chan et al., 2008; Huang et al., 2005), a portion (up to 50%) of the oxycodone may have been cleared from the circulation. However, it is possible that this delay in testing had no effect on the behavior of the rats in the small open field. Although the time course of the locomotor effects of oxycodone have not been investigated, the antinociceptive effects of oxycodone (IP) are stable for about 60 min (Holtman Jr and Wala, 2006). More importantly, given the close temporal proximity of the small open field and elevated plus maze test, the absence of oxycodone effects in the small open field indicates that the oxycodone effects on elevated plus maze measures were not secondary to effects on locomotor activity.

We are not aware of prior studies that have investigated the effects of oxycodone in the elevated plus-maze test; however, the effects of other opioids have been investigated in this task. Morphine administration in male Wistar rats increases open arm entries and time on the open arms in the elevated plus-maze test but does not significantly affect the percentage of open arm entries (Koks et al., 1999). Furthermore, Rezayof and associates showed that morphine increases the percentage of open arm time and percentage open arm entries in male Wistar rats without affecting locomotor activity (Rezayof et al., 2009). It has also been reported that chronic administration of fentanyl (14 days) in male C57BL/6J mice decreases anxiety-like behavior (Fujii et al., 2019). The selective mu-opioid receptor agonist DAMGO did not affect anxiety-like behavior in the elevated plus-maze test in Wistar rats (Sudakov et al., 2015) or adult C57BL/6J male mice (Kudryavtseva et al., 2004). In contrast, stimulation of delta-opioid receptors increases the percentage time on the open arms and open arm entries in male Lewis rats in the elevated plus-maze test (Saitoh et al., 2004). Overall, these findings indicate that widely used opioid analgesics (oxycodone, morphine, and fentanyl) decrease anxiety-like behavior in the elevated plus-maze test. These prescription opioids are mu-opioid receptor agonists but may also act via delta and kappa-opioid receptors (Kristensen et al., 1994; Maguire et al., 1992; Nielsen et al., 2007; Yang et al., 2016). It is currently not known via which opioid receptor oxycodone exerts its anxiolytic effects.

In the present study, we also observed sex differences in the elevated plus-maze test and the small open field test. In the small open field test, the females traveled a greater distance, had more horizontal beam breaks, and displayed more rearing. In prior studies, we investigated sex differences in Wistar rats in the small open field test, but sex differences were not observed for all activity parameters (Bruijnzeel et al., 2016; Chellian et al., 2021; Chellian et al., 2020). This is most likely due to the fact that the sex differences are relatively small, and there is significant variability between animals in activity parameters. Overall, relatively large group sizes and high statistical power (0.99 for the present study) are needed to detect sex differences in animal studies (Button et al., 2013). In the elevated plus-maze test, the females had a higher percentage of open arm entries and spent a higher percentage of time on the open arms. The females also made more open arm entries, spent more time on the open arms, spent less time in the closed arms, and traveled a greater distance than the males. The number of closed arm entries was not affected by the sex of the rats, thus indicating that the males were as active as the females in the relatively protected areas of the maze. Overall, these findings are in line with our previous studies in which we found that females display less anxiety-like behavior in the elevated plus-maze test (Chellian et al., 2021; Chellian et al., 2020; Knight et al., 2021). This sex difference in anxiety-like behavior might be explained by sex differences in hormone levels during development. Male rats display less anxiety-like behavior when they are castrated before puberty (Domonkos et al., 2017). Treatment with testosterone does not reverse the long-term effects of hypogonadism on anxiety-like behavior in adult males (Domonkos et al., 2017). Another study reported that pre-pubertal castration increases locomotor activity in the open field and elevated plus-maze during adolescence and in adulthood (Renczés et al., 2020). These findings suggest that the relatively low level of activity and a high level of anxiety-like behavior in the males might be due to testosterone exposure during development (Domonkos et al., 2017; Domonkos et al., 2018).

In conclusion, the present findings indicate that oxycodone induces a large decrease in anxiety-like behavior in the elevated plus-maze test and that oxycodone had a greater anxiolytic effect in males than females. In the elevated plus-maze test, the females displayed less anxiety-like behavior than the males, and the females were more active in the elevated plus-maze test and small open field test.

## CRediT authorship contribution statement

**A. Bruijnzeel:** Conceptualization, Formal analysis, Writing - Original Draft, Visualization, Supervision, Funding acquisition. **A. Behnood-Rod:** Investigation, Project administration. **W. Malphurs:** Investigation. **R. Chellian:** Formal analysis, Investigation, Writing - Review & Editing, Visualization. **Robert M. Caudle:** Conceptualization, Funding acquisition. **M. Febo:** Conceptualization, Writing - Review & Editing. **B. Setlow:** Conceptualization, Writing - Review & Editing, Funding acquisition. **J. Neubert:** Conceptualization, Writing - Review & Editing, Project administration, Funding acquisition.

## REFERENCES

Adamec, R.E., Sayin, U., Brown, A., 1991. The effects of corticotrophin releasing factor (CRF) and handling stress on behavior in the elevated plus-maze test of anxiety. J.Psychopharmacol. 5(3), 175–186.

Alexeeva, E., Nazarova, G., Sudakov, S., 2012. Effects of Peripheral μ, δ, and κ-Opioid Receptor Agonists on the Levels of Anxiety and Motor Activity of Rats. Bulletin of experimental biology and medicine 153(5), 720–721.

Boyd, C.J., McCabe, S.E., Cranford, J.A., Young, A., 2006. Adolescents’ motivations to abuse prescription medications. Pediatrics 118(6), 2472–2480.

Bruijnzeel, A.W., Lewis, B., Bajpai, L.K., Morey, T.E., Dennis, D.M., Gold, M., 2006. Severe deficit in brain reward function associated with fentanyl withdrawal in rats. Biol.Psychiatry 59(5), 477–480.

Bruijnzeel, A.W., Qi, X., Guzhva, L.V., Wall, S., Deng, J.V., Gold, M.S., Febo, M., Setlow, B., 2016. Behavioral Characterization of the Effects of Cannabis Smoke and Anandamide in Rats. PloS one 11(4), e0153327.

Button, K.S., Ioannidis, J.P., Mokrysz, C., Nosek, B.A., Flint, J., Robinson, E.S., Munafò, M.R., 2013. Power failure: why small sample size undermines the reliability of neuroscience. Nature reviews neuroscience 14(5), 365–376.

Centers for Disease Control and Prevention, N.C.f.H.S., 2021. Multiple Cause of Death 1999-2019, CDC WONDER Online Database, released 12/2020. (10/18/2021).

Chan, S., Edwards, S.R., Wyse, B.D., Smith, M.T., 2008. Sex differences in the pharmacokinetics, oxidative metabolism and oral bioavailability of oxycodone in the Sprague-Dawley rat. Clinical and Experimental Pharmacology and Physiology 35(3), 295–302.

Chellian, R., Behnood-Rod, A., Wilson, R., Febo, M., Bruijnzeel, A.W., 2021. Adolescent nicotine treatment causes robust locomotor sensitization during adolescence but impedes the spontaneous acquisition of nicotine intake in adult female Wistar rats. Pharmacology Biochemistry and Behavior 207, 173224.

Chellian, R., Behnood-Rod, A., Wilson, R., Wilks, I., Knight, P., Febo, M., Bruijnzeel, A.W., 2020. Exposure to smoke from high-but not low-nicotine cigarettes leads to signs of dependence in male rats and potentiates the effects of nicotine in female rats. Pharmacology Biochemistry and Behavior, 172998.

Chen, Q., Larochelle, M.R., Weaver, D.T., Lietz, A.P., Mueller, P.P., Mercaldo, S., Wakeman, S.E., Freedberg, K.A., Raphel, T.J., Knudsen, A.B., 2019. Prevention of prescription opioid misuse and projected overdose deaths in the United States. JAMA network open 2(2), e187621–e187621.

Collins, D., Reed, B., Zhang, Y., Kreek, M.J., 2016. Sex differences in responsiveness to the prescription opioid oxycodone in mice. Pharmacology Biochemistry and Behavior 148, 99–105.

Craft, R.M., 2008. Sex differences in analgesic, reinforcing, discriminative, and motoric effects of opioids. Experimental and clinical psychopharmacology 16(5), 376.

Domonkos, E., Borbélyová, V., Csongová, M., Bosý, M., Kačmárová, M., Ostatníková, D., Hodosy, J., Celec, P., 2017. Sex differences and sex hormones in anxiety-like behavior of aging rats. Hormones and behavior 93, 159–165.

Domonkos, E., Hodosy, J., Ostatníková, D., Celec, P., 2018. On the role of testosterone in anxiety-like behavior across life in experimental rodents. Frontiers in endocrinology 9, 441.

Febo, M., Gonzalez-Rodriguez, L.A., Capo-Ramos, D.E., Gonzalez-Segarra, N.Y., Segarra, A.C., 2003. Estrogen-dependent alterations in D2/D3-induced G protein activation in cocaine-sensitized female rats. Journal of neurochemistry 86(2), 405–412.

Feingold, D., Brill, S., Goor-Aryeh, I., Delayahu, Y., Lev-Ran, S., 2017. Misuse of prescription opioids among chronic pain patients suffering from anxiety: A cross-sectional analysis. General hospital psychiatry 47, 36–42.

File, S.E., 1990. One-trial tolerance to the anxiolytic effects of chlordiazepoxide in the plus-maze. Psychopharmacology 100(2), 281–282.

Fujii, K., Koshidaka, Y., Adachi, M., Takao, K., 2019. Effects of chronic fentanyl administration on behavioral characteristics of mice. Neuropsychopharmacology reports 39(1), 17–35.

Gros, D.F., Milanak, M.E., Brady, K.T., Back, S.E., 2013. Frequency and severity of comorbid mood and anxiety disorders in prescription opioid dependence. The American journal on addictions 22(3), 261–265.

Guimarães, F.S., Chiaretti, T., Graeff, F., Zuardi, A., 1990. Antianxiety effect of cannabidiol in the elevated plus-maze. Psychopharmacology 100(4), 558–559.

Han, B., Compton, W.M., Blanco, C., Crane, E., Lee, J., Jones, C.M., 2017. Prescription opioid use, misuse, and use disorders in US adults: 2015 National Survey on Drug Use and Health. Annals of internal medicine 167(5), 293–301.

Hayashi, M.L., Rao, B.S., Seo, J.-S., Choi, H.-S., Dolan, B.M., Choi, S.-Y., Chattarji, S., Tonegawa, S., 2007. Inhibition of p21-activated kinase rescues symptoms of fragile X syndrome in mice. Proceedings of the national academy of sciences 104(27), 11489–11494.

Hogg, S., 1996. A review of the validity and variability of the elevated plus-maze as an animal model of anxiety. Pharmacology, biochemistry, and behavior 54(1), 21–30.

Holtman Jr, J.R., Wala, E.P., 2006. Characterization of the antinociceptive effect of oxycodone in male and female rats. Pharmacology Biochemistry and Behavior 83(1), 100–108.

Huang, L., Edwards, S.R., Smith, M.T., 2005. Comparison of the pharmacokinetics of oxycodone and noroxycodone in male dark agouti and Sprague–Dawley rats: influence of streptozotocin-induced diabetes. Pharmaceutical research 22(9), 1489–1498.

Kane, S., 2021. Oxycodone, ClinCalc DrugStats Database, Version 2021.10. ClinCalc: https://clincalc.com/DrugStats/Drugs/Oxycodone. Updated September 15, 2021. Accessed October 25, 2021.

Knight, P., Chellian, R., Wilson, R., Behnood-Rod, A., Panunzio, S., Bruijnzeel, A.W., 2021. Sex differences in the elevated plus-maze test and large open field test in adult Wistar rats. Pharmacology Biochemistry and Behavior 204, 173168.

Koks, S., Soosaar, A., Voikar, V., Bourin, M., Vasar, E., 1999. BOC-CCK-4, CCKBreceptor agonist, antagonizes anxiolytic-like action of morphine in elevated plus-maze. Neuropeptides 33(1), 63–69.

Korte, S.M., De Boer, S.F., Bohus, B., 1999. Fear-potentiation in the elevated plus-maze test depends on stressor controllability and fear conditioning. Stress 3(1), 27–40.

Kristensen, K., Christensen, C.B., Christrup, L.L., 1994. The mu1, mu2, delta, kappa opioid receptor binding profiles of methadone stereoisomers and morphine. Life sciences 56(2), 45–50.

Kudryavtseva, N.N., Gerrits, M.A., Avgustinovich, D.F., Tenditnik, M.V., van Ree, J.M., 2004. Modulation of anxiety-related behaviors by mu- and kappa-opioid receptor agonists depends on the social status of mice. Peptides 25(8), 1355–1363.

Liu, Y.-L., Yan, L.-D., Zhou, P.-L., Wu, C.-F., Gong, Z.-H., 2009. Levo-tetrahydropalmatine attenuates oxycodone-induced conditioned place preference in rats. European journal of pharmacology 602(2-3), 321–327.

Maguire, P., Tsai, N., Kamal, J., Cometta-Morini, C., Upton, C., Loew, G., 1992. Pharmacological profiles of fentanyl analogs at μ, δ and κ opiate receptors. European journal of pharmacology 213(2), 219–225.

Martins, S.S., Fenton, M.C., Keyes, K.M., Blanco, C., Zhu, H., Storr, C.L., 2012. Mood and anxiety disorders and their association with non-medical prescription opioid use and prescription opioid-use disorder: longitudinal evidence from the National Epidemiologic Study on Alcohol and Related Conditions. Psychological medicine 42(6), 1261–1272.

Martins, S.S., Keyes, K.M., Storr, C.L., Zhu, H., Chilcoat, H.D., 2009a. Pathways between nonmedical opioid use/dependence and psychiatric disorders: results from the National Epidemiologic Survey on Alcohol and Related Conditions. Drug and alcohol dependence 103(1-2), 16–24.

Martins, S.S., Storr, C.L., Zhu, H., Chilcoat, H.D., 2009b. Correlates of extramedical use of OxyContin® versus other analgesic opioids among the US general population. Drug and Alcohol Dependence 99(1-3), 58–67.

Monory, K., Greiner, E., Sartania, N., Sallai, L., Pouille, Y., Schmidhammer, H., Hanoune, J., Borsodi, A., 1999. Opioid binding profiles of new hydrazone, oxime, carbazone and semicarbazone derivatives of 14-alkoxymorphinans. Life sciences 64(22), 2011–2020.

Nielsen, C.K., Ross, F.B., Lotfipour, S., Saini, K.S., Edwards, S.R., Smith, M.T., 2007. Oxycodone and morphine have distinctly different pharmacological profiles: radioligand binding and behavioural studies in two rat models of neuropathic pain. Pain 132(3), 289–300.

Pellow, S., File, S.E., 1986. Anxiolytic and anxiogenic drug effects on exploratory activity in an elevated plus-maze: a novel test of anxiety in the rat. Pharmacology Biochemistry and Behavior 24(3), 525–529.

Pradhan, A.A.A., Yu, X.H., Laird, J.M., 2010. Modality of hyperalgesia tested, not type of nerve damage, predicts pharmacological sensitivity in rat models of neuropathic pain. European Journal of Pain 14(5), 503–509.

Qi, X., Guzhva, L., Yang, Z., Febo, M., Shan, Z., Wang, K.K., Bruijnzeel, A.W., 2016. Overexpression of CRF in the BNST diminishes dysphoria but not anxiety-like behavior in nicotine withdrawing rats. Eur Neuropsychopharmacol 26, 1378–1389.

Renczés, E., Borbélyová, V., Keresztesová, L., Ostatníková, D., Celec, P., Hodosy, J., 2020. The age-dependent effect of pre-pubertal castration on anxiety-like behaviour in male rats. Andrologia 52(7), e13649.

Rezayof, A., Hosseini, S.-s., Zarrindast, M.-R., 2009. Effects of morphine on rat behaviour in the elevated plus maze: the role of central amygdala dopamine receptors. Behavioural brain research 202(2), 171–178.

Riley, J., Eisenberg, E., Müller-Schwefe, G., Drewes, A.M., Arendt-Nielsen, L., 2008. Oxycodone: a review of its use in the management of pain. Current medical research and opinion 24(1), 175–192.

Saitoh, A., Kimura, Y., Suzuki, T., Kawai, K., Nagase, H., Kamei, J., 2004. Potential anxiolytic and antidepressant-like activities of SNC80, a selective δ-opioid agonist, in behavioral models in rodents. Journal of pharmacological sciences 95(3), 374–380.

Seth, P., Rudd, R.A., Noonan, R.K., Haegerich, T.M., 2018. Quantifying the epidemic of prescription opioid overdose deaths. American Public Health Association.

Srivastava, A.B., Mariani, J.J., Levin, F.R., 2020. New directions in the treatment of opioid withdrawal. The Lancet 395(10241), 1938–1948.

Stock, H., Foradori, C., Ford, K., Wilson, M.A., 2000. A lack of tolerance to the anxiolytic effects of diazepam on the plus-maze: comparison of male and female rats. Psychopharmacology 147(4), 362–370.

Sudakov, S., Nazarova, G., Alekseeva, E., Kolpakov, A., 2015. Peripheral administration of a μ-opioid receptor agonist DAMGO suppresses the anxiolytic and stimulatory effects of caffeine. Bulletin of experimental biology and medicine 158(3), 295–297.

Sui-Hui, L., Yu-HuiΔ, L., Xiao-Wei, Y., Dong-Yan, L., Yan, C., 2013. Oxycodone hydrochloride improves the anxiety and depression in patients with moderate and severe cancer pain. Chinese Journal of Pain Medicine 7.

Sullivan, M., Edlund, M., Zhang, L., Unutzer, J., 2006. Depression and anxiety disorders predict increased use of prescribed opioids for non-cancer pain. The Journal of Pain 7(4), S50.

Tan, S., Xue, S., Behnood-Rod, A., Chellian, R., Wilson, R., Knight, P., Panunzio, S., Lyons, H., Febo, M., Bruijnzeel, A.W., 2019. Sex differences in the reward deficit and somatic signs associated with precipitated nicotine withdrawal in rats. Neuropharmacology 160, 107756.

Walf, A.A., Frye, C.A., 2007. The use of the elevated plus maze as an assay of anxiety-related behavior in rodents. Nat.Protoc. 2(2), 322–328.

Yang, P.-P., Yeh, G.-C., Yeh, T.-K., Xi, J., Loh, H.H., Law, P.-Y., Tao, P.-L., 2016. Activation of delta-opioid receptor contributes to the antinociceptive effect of oxycodone in mice. Pharmacological research 111, 867–876.

Zacny, J.P., Gutierrez, S., Kirulus, K., McCracken, S.G., 2011. Psychopharmacological effects of oxycodone in volunteers with and without generalized anxiety disorder. Experimental and clinical psychopharmacology 19(2), 85.

